# Autoantibody hotspots reveal origin and impact of immunogenic XIST ribonucleoprotein complex

**DOI:** 10.1101/2025.01.16.633465

**Authors:** Bingyu Yan, Jinwoo Lee, Suhas Srinivasan, Quanming Shi, Diana R. Dou, Srijana Davuluri, Swarna Nandyala, Adrianne Woods, Gwendolyn Leatherman, Yanding Zhao, Roman E. Reggiardo, Manasi Sawant, Hawa R. Thiam, Ami A. Shah, David F. Fiorentino, Lorinda S. Chung, Howard Y. Chang

## Abstract

Four out of five patients with autoimmune diseases are women. The XIST ribonucleoprotein (RNP) complex, comprising the female-specific long noncoding RNA XIST and over 100 associated proteins, may drive several autoimmune diseases that disproportionately affect women, who have elevated levels of autoantibodies against the XIST RNP. However, the structural distribution, potential origin, and clinical significance of XIST RNP autoantibodies remained unexplored. Here, we find that XIST RNP is associated with autoantigens associated with six female-biased autoimmune conditions. Mapping autoantibody targets to their occupancy sites on XIST shows that these autoantigens are concentrated at discrete “hotspots” along the XIST lncRNA, notably the A-repeat. Cell type-specific protein expression data nominated neutrophils as a predominant source of hotspot antigens, and we confirmed the presence of both XIST and hotspot antigens in neutrophil extracellular traps during NETosis, an immunogenic programmed cell death pathway triggered by neutrophil activation upon which neutrophils extrude their nuclear content. Furthermore, we found that levels of autoantibodies against a top hotspot antigen, SPEN, that binds the A-repeat, correlate with severe digital ischemia in systemic sclerosis in two independent cohorts. Together, these data show a plausible mechanism for the origin of AXA, guided by RNA structure and RNA-protein interactions, and show that antibodies to XIST RNP holds promise for disease endotyping and prognostication in female-biased autoimmune conditions.

**One Sentence Summary:** Novel autoantibodies target hotspots on XIST ribonucleoprotein complex in female-biased autoimmune diseases.

## INTRODUCTION

Autoimmunity exhibits a striking sex bias, with conditions such as lupus, scleroderma, and Sjögren syndrome affecting women at rates up to ten times higher than men(*1*). Multiple factors contribute to the emergence of female-predominant autoimmune diseases, including sex hormones and the escape of immunoregulatory genes from the inactivated X chromosome which is reviewed here(*1*). We have implicated the X-inactive specific transcript (XIST)—a long non-coding RNA expressed only in women and responsible for X chromosome inactivation—as a dominant driver of female-biased autoimmunity(*2*). Specifically, we found that expressing XIST in male mice is sufficient to reproduce female-level autoimmune pathology, resulting in elevated levels of autoantibodies against XIST ribonucleoproteins (RNPs), a phenomenon also observed in human patients with autoimmune diseases. However, the role of the XIST RNP complex in the etiopathogenesis of autoimmune disease remains incompletely explored.

The XIST RNP complex comprises the 19-kb XIST lncRNA, which scaffolds up to 150 XIST-associated proteins (XAPs) in different cell types to form a higher-order structure mediated by multivalent RNA-RNA, RNA-protein, and protein-protein interactions. However, how the XIST RNP complex acts as the target of autoantibodies and the immunogenicity of each XAP within the distinct functional domains of the XIST RNA remain unknown. We hypothesized that due to the differential autoantibody accessibility against the XIST RNP complex for immune recognition, certain XIST domain-specific XAPs would have more immunogenic potential than others and be associated with autoimmunity.

Here, by utilizing the Human Autoantigen Atlas (AAgAtlas)(*3*) and previous data regarding the secondary structure of XIST, as well as RNA immunoprecipitation and cross-linking immunoprecipitation sequencing data(*4*) regarding XIST-protein and protein-protein interactions, our aim was to confirm that the structural intricacies of the XIST RNP complex are directly related to immunogenicity. Furthermore, we sought to identify whether “hotspots” of autoantibody targets could provide insights into the development of autoimmune responses, and whether autoantibodies targeting these “hotspots” could have clinical correlates in sex-biased autoimmune diseases.

## RESULTS

### Autoantigens in female-biased autoimmune diseases are enriched for XIST-associated proteins

To survey the association of XIST-associated proteins (XAPs) with autoimmune diseases, we mined the human autoantigen atlas (AAgAtlas) database(*3*). In 35/104(33.65%) autoimmune diseases from the AAgAtlas, 57 of 149 (38.26%) XIST-associated proteins (XAPs) are identified as autoantigens **(Fig. 1A-C)**. 15 of 35 autoimmune diseases are observed with multiple (two or more) XAP autoantigens. Systemic lupus erythematosus (SLE) and systemic sclerosis (SSc) was reproduced to be associated with multiple AXAs, validating the study of Dou et al.(*2*); in addition, mixed connective tissue disease, multiple sclerosis, rheumatoid arthritis, inflammatory bowel disease are identified to be associated with AXA for the first time **(Fig. 1D)**. Among all XAPs, SSB, HMGB1, and HMGB2 are the top 3 autoantigens that present in 13 autoimmune diseases **Fig. 1E)**. Interestingly, we found autoantigens are enriched in XAPs in autoimmune and inflammatory diseases, such as SSc, SLE, rheumatoid arthritis (RA), and mixed connective tissue disease (p< 0.05, hypergeometric test); these autoimmune diseases have higher female-to-male ratios than the other autoimmune diseases in which the associated autoantigens are not enriched in XAPs (p < 0.01, Mann-Whitney’s U test, **Fig. 1E**). These results suggest that XAPs are major sources of autoantigens in multiple autoimmune diseases.

**Fig. 1.**
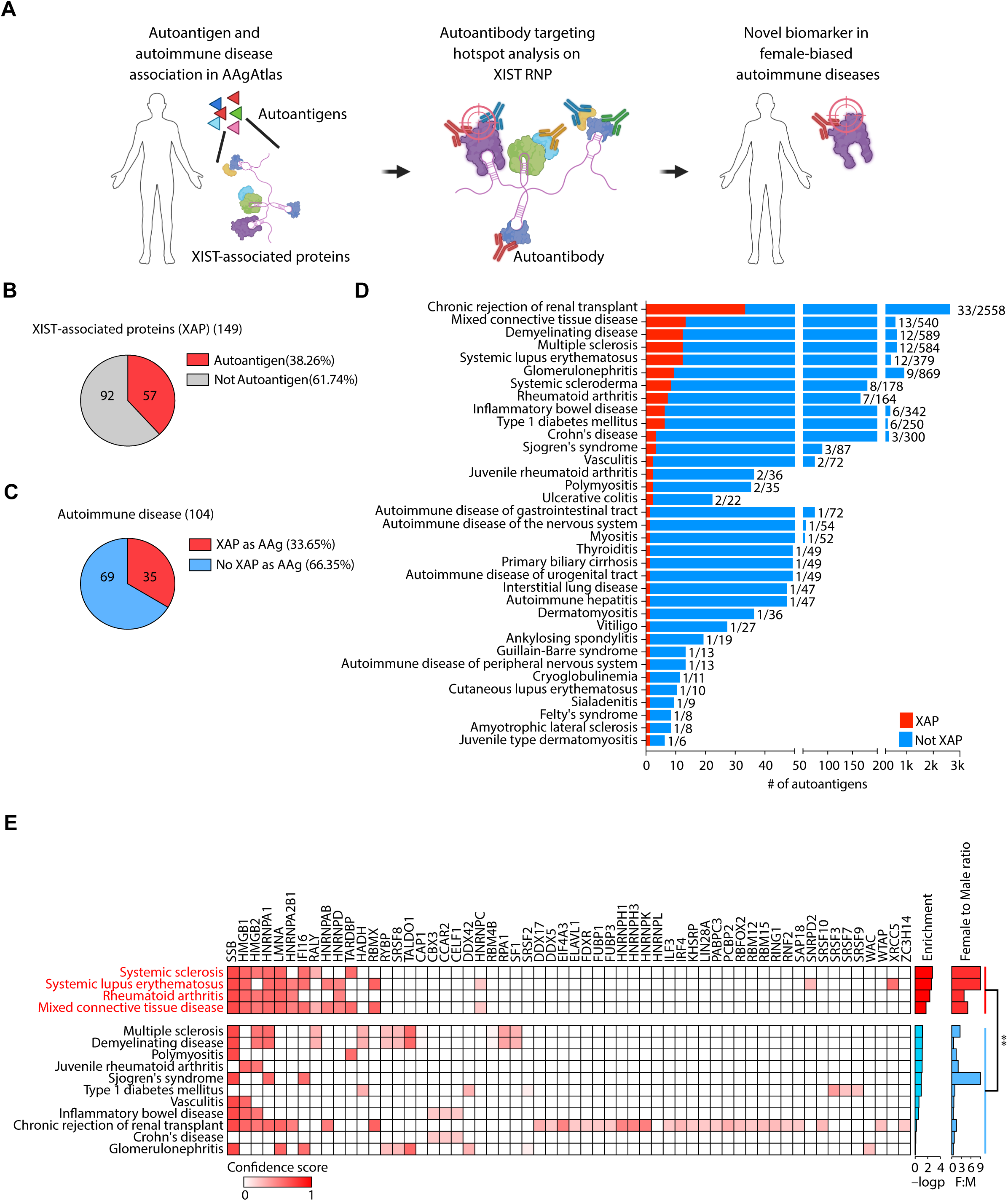
Autoantigens in female-biased autoimmune diseases are enriched for XIST-associated proteins. (**A**) Flowchart for this study. (**B**) Numbers of XIST-associated proteins (XAPs) that are autoantigens among all XAP. (**C**) Numbers of autoimmune diseases that have XAP as autoantigens among all autoimmune diseases in the Human Autoantigen Atlas. (**D**) Number of autoantigens in autoimmune diseases. XAP are highlighted in red. (**E**) The association between antigenic XAP and autoimmune diseases. Association confidence scores obtained from (*3*) are displayed as heatmap. -log10(p-value) from hypergeometric tests are displayed as bar graphs with *p* < 0.05 as red bars. Female to male ratios for each autoimmune disease are displayed as bar graph, red bars indicate autoantigens in these diseases are enriched in XAP, blue bars indicate autoantigens in these diseases are not enriched in XAP; \*\**p* < 0.01, Mann-Whitney U test.

### Immunogenicity of XAPs is associated with XIST functional domains

The XIST ribonucleoprotein complex forms a higher-order structure mediated by RNA-RNA, RNA-protein, and protein-protein interactions which project differential accessibility of autoantigens to immunorecognition. We used a three-step process to generate a comprehensive map of XIST RNP and visualize its immunogenicity **(Fig. 2A-C)**. First, we visualized the XIST RNA secondary structure as a function of its primary sequence. XIST consists of a series of repeats, termed A-F, and XIST RNA can base pair with itself across short and long distances in a hierarchical fashion. XIST secondary structure was determined by psoralen crosslinking in living cells and deep sequencing(*4*); each arc on top of Fig. 2C represents an RNA duplex along XIST RNA. Second, we mapped the XAPs curated across multiple cell types to their occupancy on XIST RNA as defined by published formaldehyde RNA immunoprecipitation and sequencing (fRIP-seq) or enhanced crosslinking immunoprecipitation (CLIP-seq)(*4*)(see methods). After mapping the XAP occupancy on XIST RNA, we performed a k-means clustering based on the fRIP-seq or eCLIP-seq signal which reflects the binding probability of each protein to the RNA and grouped the autoantigens to XIST functional domains **(Fig. 2C)**. This visualization revealed the modular architecture of XIST RNP; specific XAPs occupied the A, F, BCD, or E domains of XIST that reflected the XIST RNA secondary structure, concordant with prior studies. Third, to evaluate the immunogenicity of XAPs, we visualized the autoantibody frequencies to each XAP across 125 individuals from the general population and autoimmune patients (with, SLE, SSc, or dermatomyositis) **(Fig. 2D)**. Autoantibody reactivity is projected onto the XIST RNA and ranked by the number of samples which have AXA reactivity with MFI>100 (a proxy for autoantibody titer) from top to the bottom in each functional domain.

**Fig. 2.**
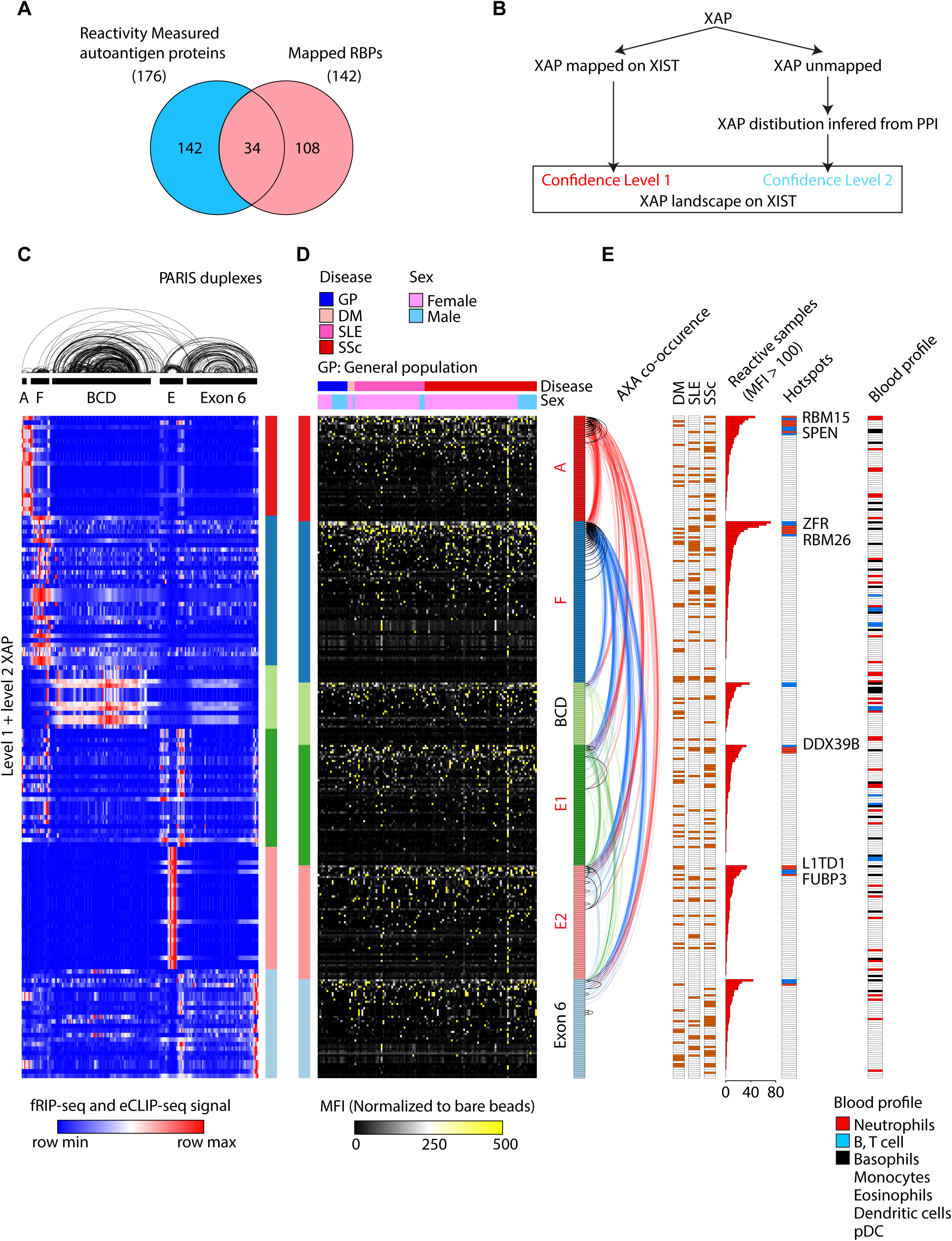
Immunogenicity of XAPs is associated with XIST functional domains. (**A**) Overlap of reactivity quantified antigenic proteins with XIST-associated proteins with available location information. Number of reactivity-quantified antigenic proteins with available location information, displayed as intersection from the Venn diagram. **(B)** Workflow for assigning and inferring the location of antigenic proteins in XIST RNA by protein-protein interaction. (**C**) Binding profile for XIST-associated proteins (XAP). Row normalized binding intensity for each XAP are displayed as heatmap in each row within 100-bp windows across XIST RNA. XIST secondary structure was determined by psoralen crosslinking in living cells and deep sequencing(*4*); each arc on top represents a RNA duplex along XIST RNA. Clusters of XAP are indicated by the bar on the right-hand side. **(D)** Reactivity of XAP derived autoantigens across general population and autoimmune disease patients. Disease annotations and sex annotations are on the top of the heatmap. Annotations for XAP-derived autoantigens in different XIST functional domains are shown on two sides of the heatmap. (**E**) Functional annotations for XAP-derived autoantigens. Panels from left to right are, 1) Pairwise co-occurrence of autoantigens (black curves indicate intra-domain co-occurrence, colored curves indicate inter-domain co-occurrence with color assignment established from the upper domain into another domain); 2) Significantly elevated autoantigens in any autoimmune disease from(*2*); 3) Numbers of samples that have MFI > 100 reactivity; 4) Immunogenic hotspots: hotspots that are significant elevated in any autoimmune disease are shown in red, others are shown in blue; 5) Blood profile annotation from the human protein atlas; 6) Immunogenic hotspots in pristane-induced SLE mouse samples.

Our map immediately revealed that autoantibodies to XIST RNP are not randomly distributed. Certain regions of the XIST RNP elicited high titer antibodies across many more patients than other regions of the XIST RNP; we operationally defined the top 10% most prevalent AXA as AXA hotspots **(Fig. 2E)**. Importantly, repeats A, E and F are coordinately occupied by AXA hotspots that are significantly elevated in autoimmune disease patients, and co-occurrence of autoantigens along XIST is more clustered than expected by chance alone (p < 0.01 for within A repeat or F repeat or inter A-F repeats, Fisher exact test, **Fig. 2E**). The A-repeat is significantly associated with greater frequency of highly reactive AXAs (Odds ratio = 2.55); the F-repeat is the second domain with most prominent hotspots (Odds ratio = 0.94). Two additional focal hotspots around E1 and E2 regions are also identified. These results suggest that XIST is an RNA scaffold that concentrates specific antigenic proteins and displays them in a polymeric fashion and the AXA hotspots in XIST repeat A and F regions are shared more frequently among autoimmune patients.

### NETosis releases XIST-associated proteins to cell-free environment

Why are some XAPs more immunogenic than others? In addition to the portion of XIST that they occupy, we wondered whether XAPs released from specific cell types can be more reactive. As our list of XAP is curated from multiple cell types, we deconvoluted the list using data available from the Human Protein Atlas: <u>Human Protein Atlas</u> proteinatlas.org(*5*). Most of the XAPs in hotspots are highly expressed and enriched in neutrophils; several of the hotspot XAPs are also expressed in B cells. Many of the less immunogenic XAPs are broadly expressed across blood cell types **(Fig. 2E)**. These results suggest that in addition to the XIST RNA scaffold, their release from specific cell types may also contribute to their immunogenicity as determined by autoantibodies.

XIST RNP is localized to the condensed inactive X chromosome in the nucleus, termed the Barr body, and XIST RNP must somehow be exposed to the extracellular space to elicit antibodies. Neutrophils release their nuclear content to the extracellular environment in the form of neutrophil extracellular traps (NETs), which are web-like structures composed of DNA, histones, and antimicrobial proteins during a specialized form of cell death called NETosis(*6*). A recent study indicated that neutrophils are the main source of cell-free DNA in cancer patients(*7*). Interestingly, we observed neutrophils as the dominant cell source for autoantigens in our panel with known blood profile information (47.83%, **Fig. 3A**) from the Human Protein Atlas (<u>Human Protein Atlas</u> proteinatlas.org) (*5*). Among the 22 autoantigens that are inferred to be coming from the neutrophil, we ranked them by prevalence in our cohort **(Fig. 3B)** and hypothesized that these autoantigens are released from neutrophils during NETosis.

**Fig. 3.**
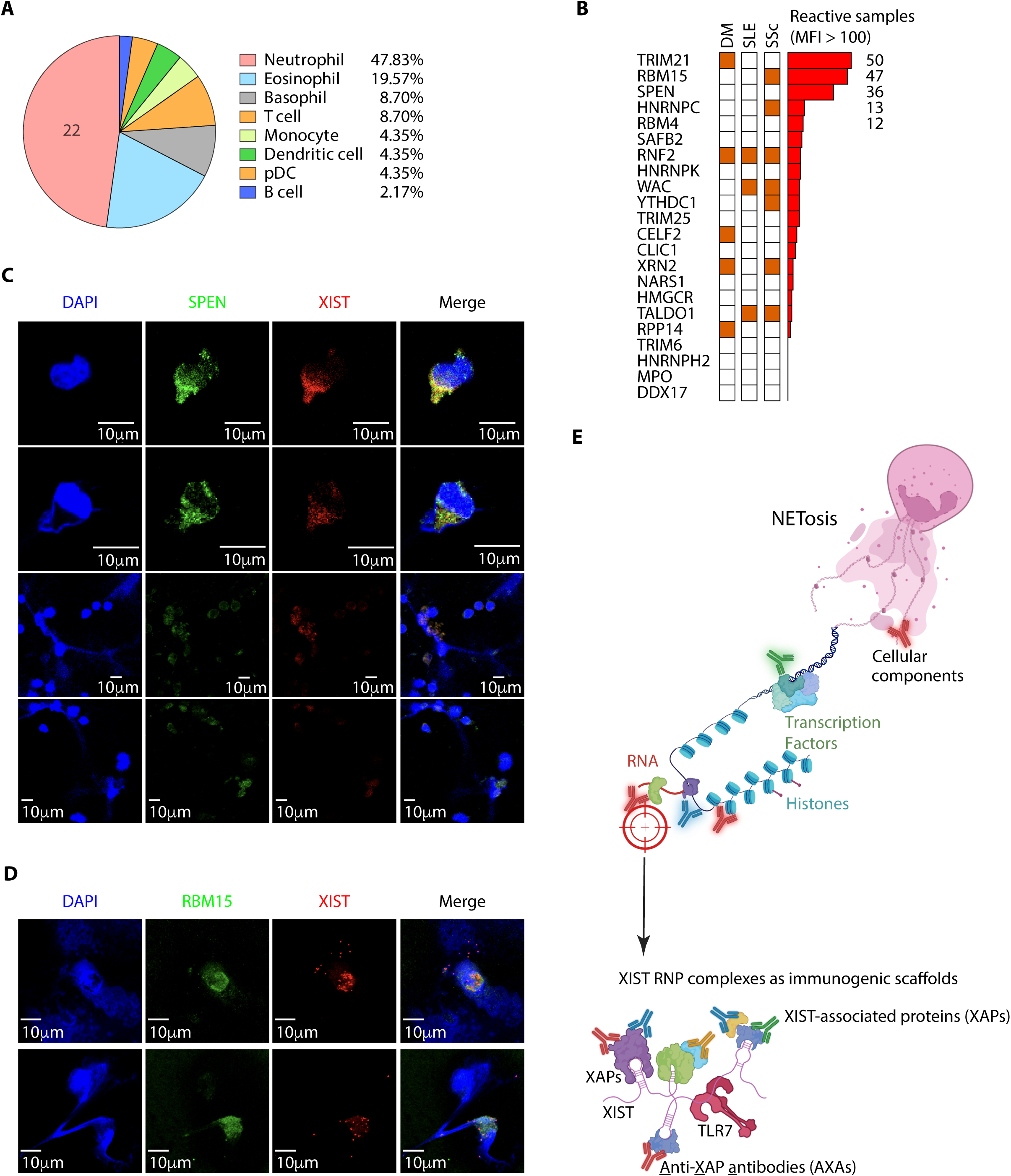
NETosis releases XIST-associated proteins to a cell-free environment. (**A**) Cell of origin for antigenic proteins. Proportions for each cell type are displayed as a pie chart. (**B**) Twenty-two neutrophil-associated antigenic proteins are displayed on the *y*-axis, and the degree of positive association with autoimmune disease are displayed as heatmaps. The number of human samples with MFI > 100 for each protein (sera reactivity) are shown as a bar graph. **(C)** dHL-60 cells were stimulated with ionomycin to induce NETosis and stained with DAPI (blue), anti-SPEN (green), and XIST FISH probe (red). (**D**) Human neutrophils were stimulated with PMA to induce NETosis and stained with DAPI (blue), anti-RBM15 (green), and XIST FISH probe (red). (**E**) Model of the XIST RNP complex acting as an immunogenic scaffold. Created in BioRender.

To experimentally evaluate this concept, we induced NETosis in female primary human neutrophils and performed immunofluorescence for the hotspot XAP SPEN and RBM15, which both bind the A-repeat, and concurrently performed RNA in situ hybridization for XIST RNA. Indeed, SPEN and RBM15 are released with XIST RNA during NETosis, and the XIST RNP remains bound to DNA in extracellular NETs reaching >20-30 microns in length **(Fig. 3C and D)**. The results support a model where NETosis exposes XIST RNP to cell-free environment that can be recognized by multiple autoantibodies and the RNA itself could serve as a ligand for TLR-7(*8*); this synergistic effect may promote the immune response to the XIST RNP **(Fig. 3E)**.

### SPEN autoantibody is associated with severe peripheral vascular manifestations in systemic sclerosis

Among XIST-associated protein hotspots in the A-repeat, SPEN has a rich body of literature regarding its role as a key RNA-binding protein that is required for XIST function. SPEN, also known as SHARP, is a scaffold protein where the N-terminus contains four RRM domains that directly interact with XIST A-repeat and the C-terminus contains a SPOC domain that recruits histone deacetylase and additional repressive complexes (**Fig. 4A**) (*9–11*). Significantly, SPEN has never been identified as an autoantigen in any prior studies and is not present in the Human AAgAtlas, which curates published literature. Upon review of serologic data from our previously published antigen array, which analyzed sera from female patients with systemic sclerosis (SSc) for reactivity against a panel of XAP antigens, we found that reactivity against a SPEN protein fragment **(Fig. 4A)** was detectable by fluorescent bead assay in SSc patients with a wide distribution of fluorescence signal intensities **(Fig. 4B)**. When comparing SSc patients with high and low reactivities against SPEN *(n* = 10 and *n* = 14, respectively), defined as an MFI of at least twice the median value of the distribution of fluorescence intensities in the cohort, we found that high sera reactivity against SPEN in the Stanford cohort was associated with a severe vasculopathic phenotype **(Fig. 4C)**, most significantly manifesting in the form of digital ischemia often leading to digital ulcers or digital gangrene. Because of the small size of the Stanford patient cohort, which is an inherent limitation in the investigation of rare diseases, statistical significance was only seen for the clinical outcome of digital gangrene/amputation (difference in proportion = 0.40, 95% CI = 0.09 to 0.70, *p* = 0.0088, Barnard’s exact test), though differences not reaching statistical significance could also be seen for the pathologically related outcome of digital ulcers (difference in proportion = 0.23, 95% CI = -0.16 to 0.55, *p* = 0.25, Barnard’s exact test). We sought to validate this finding in an independent cohort, with analysis performed by an independent investigator. In a larger cohort of thirty-five SSc patients from the Johns Hopkins University, we found that a statistically significant association was found between high anti-SPEN reactivity and severe Raynaud phenomenon (difference in proportion = 0.37, 95% CI = 0.047 to 0.60, *p* = 0.031, Barnard’s exact test, **Fig. 4D)**, which was defined as Raynaud phenomenon associated with digital pitting scars, digital tip ulceration, and digital gangrene **(Fig. 4E)**, in addition to a statistically significant association between high anti-SPEN antibody and digital ulceration (difference in proportion = 0.33, 95% CI = 0.0033 to 0.60, *p* = 0.044, Barnard’s exact test). These results highlight the potential of autoantibodies against hotspot XAPs such as SPEN as novel indicators of autoimmune phenotypes.

**Fig. 4.**
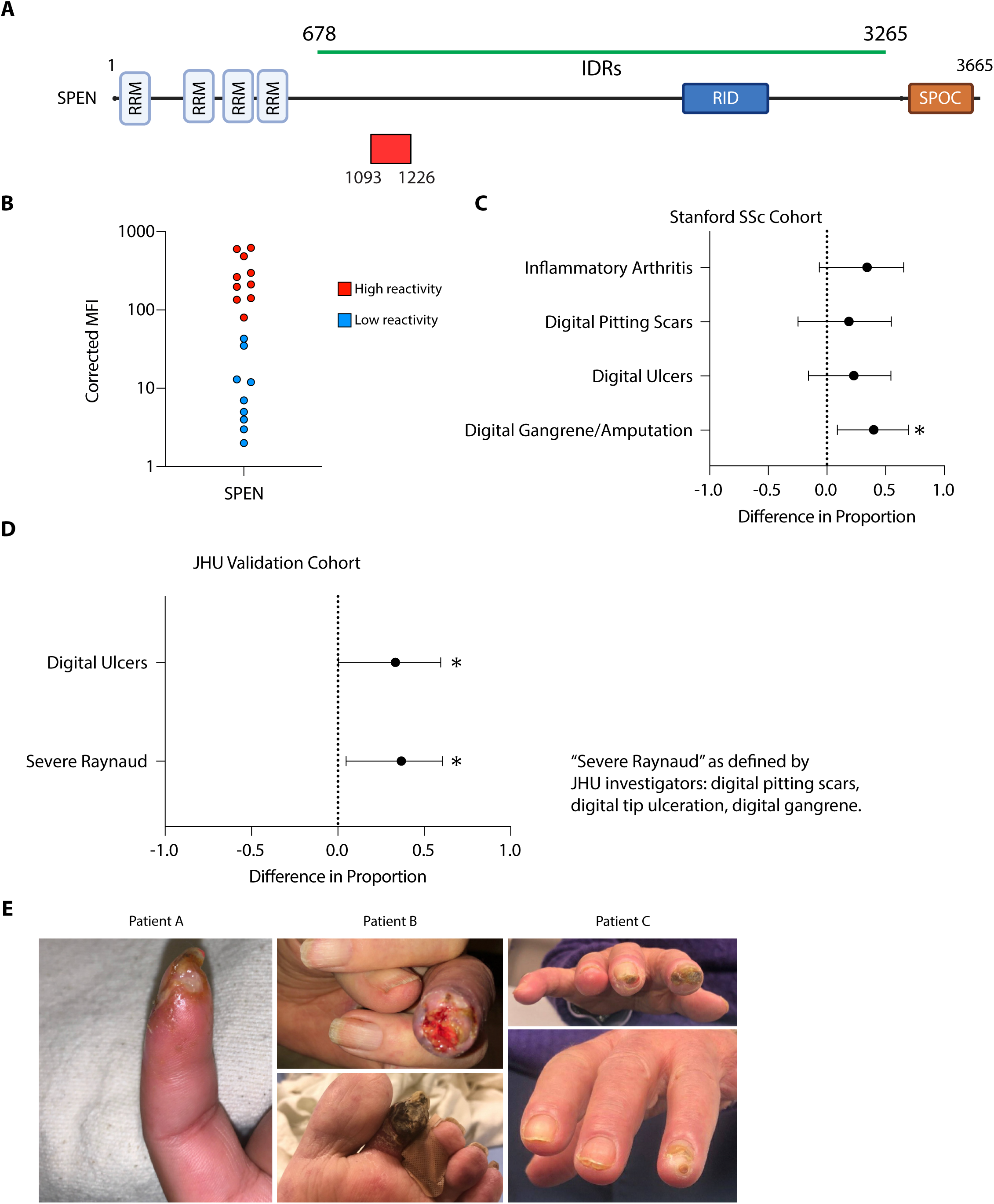
An AXA hotspot is associated with high-risk vasculopathy in systemic sclerosis patients. **(A)** Schematic showing the location of the protein fragment used in the fluorescent bead-based antigen array to assess sera reactivity against SPEN. RRM, RNA recognition motif. RID, receptor interaction domain. SPOC, Spen paralog and ortholog C-terminal. IDRs, intrinsically disordered regions. **(B)** Distribution of mean fluorescence intensity (MFI) values for sera reactivity against SPEN in the Stanford scleroderma cohort. **(C)** Calculated difference in proportion for categorical variables of interest in the Stanford scleroderma cohort as defined by Stanford investigators, with horizontal lines indicating the 95% confidence intervals, and a dotted vertical line at 1.0 signifying no association. * *p* < 0.05. **(D)** Calculated difference in proportion for digital ulcers and severe Raynaud phenomenon (*i.e.* digital pitting scars, digital tip ulceration, digital gangrene) in the JHU validation scleroderma cohort, as defined by JHU investigators. * *p* < 0.05. **(E)** Representative clinical photos of severe digital ulceration and gangrene in three patients with high anti-SPEN sera reactivity.

## DISCUSSION

Here, we demonstrated that XIST-associated proteins (XAPs) are a major source of autoantigens in many female sex-biased autoimmune diseases. Many of these autoantigens are RNA binding proteins previously documented as the targets of autoantibodies(*3*), and the recognition that these autoantibodies all target a shared RNA protein complex provides a unifying theme to previously disparate observations. The female-specific XIST ribonucleoprotein (RNP) complex provides a polymeric scaffold for antigenic proteins, and has been found to be sufficient to drive autoimmunity in mouse models of lupus, and furthermore, autoantibodies against the XIST RNP complex (AXA) can be seen in diseases such as systemic sclerosis, systemic lupus erythematosus, and dermatomyositis(*2*). By systematically analyzing where each AXA binds to on the XIST lncRNA, we find that XAP immunogenicity is concentrated in “hotspots” along the XIST sequence. **(Fig. 2E)**. The XAP hotspots were located within discrete XIST binding domains, which would be able to then dictate differential accessibility of the antigenic protein for immune recognition.

We also observed that more than one-third of autoimmune diseases are associated with XAP-derived autoantigens **(Fig. 1B)**. XAPs are major sources of autoantigens in four autoimmune diseases which have a strong female sex bias, such as systemic sclerosis (4:1), systemic lupus erythematosus (9:1), rheumatoid arthritis (4:1), and mixed connective tissue disease (5:1)(*1, 12*). Though numerical enrichment of XAP autoantigens was not observed for Sjogren syndrome, an autoimmune disease with a substantial female preponderance (ranging from 6:1 to 16:1 in studies), we note that SSB (Sjogren Syndrome type B) protein and hnRNP A1 are both XAPs and are highly associated with Sjogren syndrome. Because the enrichment analysis only takes into consideration the number of XAP autoantigens associated with an autoimmune condition, rather than the individual strength of association between each autoantibody and the disease, the analysis will underreport the significance of XAP autoantibodies in diseases with a smaller number of reported autoantibodies. Nevertheless, it is reassuring that the only sex-biased condition that was not explicitly flagged as enriched in XAP autoantibodies was a condition in which an XAP autoantibody (anti-SSB) is well-known to be associated with the condition.

The risks conferred by the presence of autoantibodies are not equal and the presence of multiple antibodies may signal different immunogenic environments, especially when considering epitope accessibility, which is associated with RNP structure and specific binding domains. SSB and HNRNPA1 are in repeat F, which is the second-ranked functional domain with AXA hotspots. HMGB1 is associated with all five female sex-biased autoimmune diseases and located in repeat A. LNMA is in repeat E which is associated with all the five female-biased autoimmune diseases. In our model of XIST RNP scaffolding autoantigens, our hotspot analysis indicated A, F and E are the top 3 repeats that harbor autoimmune disease-associated AXA hotspots **(Fig. 2E)**, and our co-occurrence analysis indicated that the A and F repeats are concentrated with hotspots as well as co-occurrence AXAs which suggests they might have synergistic effects in female-biased autoimmunity **(Fig. 2E)**. Whether subdivision of AXAs by XIST binding domains can be of clinical or mechanistic relevance will be an active area of future inquiry.

Intracellular autoantigens, such as XAPs and antigens derived from ribonucleoproteins, must be released from the cell for immune recognition, which has been hypothesized to be due to necrotic cell death. An alternative mechanism of programmed cell death called NETosis presents an enticing model for autoantigen release from neutrophils, particularly as the contents of NETs are predominantly DNA and other nuclear content(*7*). We find that the expression of many hotspot XAPs are enriched in neutrophils **(Fig. 3A**), and NETosis releases XIST RNPs in association with neutrophil DNA, where the polymeric nature of XIST, XAPs, and chromatin are preserved and spatially extended in the extracellular milieu **(Fig. 3C and D)**. These results suggest that NETosis is an active mechanism by which XIST RNP particles can be released in an immunogenic environment.

Importantly, based on our analysis of XIST-associated proteins and detailed patient phenotyping, we describe a novel autoantibody with a clinically significant association, demonstrated in two independent cohorts of patients with systemic sclerosis. SSc patients with high anti-SPEN antibodies were at significantly higher risk of suffering from severe vasculopathy as manifested by digital pitting scars, debilitating digital ulceration and gangrene, in some cases leading to amputation in the Stanford cohort of SSc patients. SPEN is a large protein (390 kD) with multiple RNA recognition motifs as well as intrinsically disordered regions (IDRs) capable of binding to and aggregating proteins (*11, 13*), suggesting that SPEN may play a key role in the generation of large RNA-protein complexes. SPEN undergoes phase separation, and high-resolution imaging showed that the intact mouse Xist RNP contains 2 molecules of Xist and approximately 50 molecules of SPEN, yielding a supramolecular complex(*13*). Interestingly, we find that the anti-SPEN antibody that our assay detects is directed against a sequence that not only lies in the region of SPEN known to contain a high density of IDRs (between the RRMs and the SPOC domain) but is also within an annotated IDR (positions 1070-1287, UniProt). We hypothesize that patients who have autoantibodies against SPEN may be at higher risk of developing large immune complexes that can lead to vasculopathy and digital ischemia. The mechanism by which anti-SPEN autoantibodies may confer an increased risk of ischemia will be an active area of future research. We will also test the utility of anti-SPEN antibodies as a prognostic biomarker for the development of digital ischemia in SSc patients.

By integrating structural data with clinical and immunological insights, this study advances our understanding of the relationship between XIST and autoimmunity, offering new avenues for early detection, monitoring, and targeted therapies in autoimmune disorders. Because the appearance of autoantibodies often occurs before the onset of clinically obvious diseases, expansion of the known pool of autoantibodies can be critical for the detection and monitoring of autoimmune diseases. Though the sensitivity, specificity, and significance of individual AXA autoantibodies will each require further clinical validation, this opens a promising targeted approach to the identification of novel autoantibodies in autoimmune conditions. Our study is limited by small patient size and a single time point for sera collection; future studies with larger cohorts of autoimmune patients and with sera from multiple time points before and during disease progression are planned to be able to more definitively confirm as well as predict clinical outcomes.

## MATERIALS AND METHODS

### Human Autoantigen Atlas data mining

All available data including autoantigen and the associated diseases were acquired from the Human Autoantigen Atlas (AAgAtlas)(*3*). Autoantigens associated with autoimmune diseases were kept. Autoimmune diseases which have XIST-associated proteins as autoantigens were exacted and the number of autoantigens were calculated. Sex ratios of autoimmune diseases were obtained from published literature shown in Supplementary Table 1.

### Protein-protein interaction analysis

Proteins comprising autoantigens were first mapped to XIST RBPs which have fRIP-seq or eCLIP-seq data by physical protein-protein interaction from multiple databases by Metascape (*14*). The proteins that comprised the remaining autoantigens and unassigned were mapped to other RBPs capable of binding XIST from the fRIP-seq or eCLIP-seq data. The interacting partner with the highest score was kept in the analysis.

With the XAPs assigned onto the XIST RNA sequence, *k*-means clustering was performed with values from fRIP-seq or eCLIP-seq data, and these clusters were assigned to XIST functional domains.

### AXA Hotspot identification from human anti-XIST autoantibody data

The number of individuals with reactivity of a particular AXA with normalized (AXA - bare beads) MFI>100 is counted. Top 10% of the autoantigens are defined as AXA hotspots in human and mouse data. All the autoantibody reactivity data were obtained from (*2*).

### Autoantigen pairwise co-occurrence analysis

To calculate the pairwise co-occurrence of autoantigens in our human donor cohort, dot product between the AXA reactivity matrix and its transposed matrix is calculated with the values in AXA reactivity matrix are converted to binary indicator with normalized MFI>100 as 1 and <100 as 0. AXA that are co-occurrent in >5% (co-occurring number of individuals/total number of human donors) individuals are counted as co-occurrent AXA.

### Human protein atlas data mining

Data were acquired from the Human Protein Atlas (www.proteinatlas.org) (*5*). The blood profile of all proteins was retrieved and matched to XIST-associated proteins to indicate the main cell types that contributed to protein expression in blood.

### Neutrophil isolation

Human neutrophils were isolated from the blood of healthy donors collected by the Research and Clinical Services team at the Stanford Blood Center or by customized collection from a commercial vendor (Creative Biolabs). When isolation was performed from whole blood in our laboratory and not from a commercial vendor, we performed density gradient-based cell separation, described as follows. In brief, 7 ml of heparinized blood was layered on 7 ml of Histopaque-1119 (Sigma-Aldrich, 11911) in a 15-ml Falcon tube, and centrifuged at 800 × *g* for 20 minutes at room temperature. The upper layer and interphase were then aspirated and discarded, while the diffuse red phase above the erythrocyte pellet was collected in a new tube, washed with HBSS, then layered onto a 10-ml density gradient consisting of successive layers of 65, 70, 75, 80, and 85% Percoll (Sigma-Aldrich, P4937) and centrifuged at 800 × *g* for 20 minutes at room temperature. The interface between the 65% and 75% Percoll layers were collected in a new 15-ml tube, washed, then resuspended in RPMI at 1.5 × 10^6^ cells per ml for neutrophil activation.

### Human neutrophil activation

Human neutrophils were stimulated by phorbol myristate acetate (PMA) as previously described(*15*). In brief, isolated neutrophils were incubated for 1-3 hours with 30 nM PMA (Sigma-Aldrich, P8319) at 37 °C, then fixed with 1% formaldehyde for 10 minutes at room temperature, and spun down onto a glass slide with Cytospin at 1000 r.p.m. for 5 minutes at room temperature for FISH and immunostaining.

### HL-60 cells

HL-60 cells (ATCC CCL-240) were cultured at 37 °C and 5% CO2 in RPMI 1640 plus L-glutamine with 25 mM HEPES buffer, 1% penicillin/streptomycin, and 15% heat-inactivated FBS and differentiated to neutrophil-like cells by addition of 1.3% of DMSO. For NETosis, HL-60 cells were stimulated with ionomycin (4 μM) for 2 h.

### FISH and immunofluorescence imaging

After fixation, slides were washed with PBS, then Stellaris™ RNA FISH Probes recognizing XIST and labelled with Quasar™ 570 dye (SMF-2038-1, LGC, Biosearch Technologies, Petaluma, CA) were hybridized to samples, following the manufacturer’s instructions available online at www.biosearchtech.com/stellarisprotocols. FISH and IF were performed according to the manufacturer-provided protocol without any significant deviations. Briefly, after allowing slides to air dry following cell permeabilization with 70% ethanol for 1 hour at 4 °C, slides were immersed in Stellaris® RNA FISH Wash Buffer A (SMF-WA1) for 2-5 minutes at room temperature. Then, slides were hybridized overnight for 16 hours in a light-protected humidified chamber at 37 °C in 125 nM of Human XIST Stellaris® FISH Probes with Quasar® 570 Dye (SMF-2038-1) hybridization buffer containing the probe and appropriately diluted primary antibody. After washing, the slides were then incubated with appropriately diluted secondary antibody in Wash Buffer A for 30 minutes in a light-protected chamber at 37 °C, followed by immersion in Stellaris® RNA FISH Wash Buffer B (SMF-WB1) for 2-5 minutes at room temperature. Slides were mounted with Vectashield Mounting Medium with DAPI (Vector Laboratories, H-1800) and a cover glass.

Slides were imaged at Stanford University - CSIF: Cell Sciences Imaging Facility, RRID:SCR_017787 on the Leica Stellaris 8 DIVE, with data acquisition using LAS X software.

Primary antibodies were used as follows: rabbit anti-SPEN (1:250, Abcam, ab72266), rabbit anti-RBM15 (1:100, Proteintech, 10587-1-AP).

Secondary antibodies we used as follows: goat anti-rabbit-AF488 (1:400, Thermo Fisher, A11008) or goat anti-mouse-AF647 (1:400, Thermo Fisher, A-21244), goat anti-mouse-AF488 (1:400, Thermo Fisher, A-32723).

### Clinical cohorts

All human patient samples were de-identified in this study and obtained under the respective institutions’ IRB-approved protocols. Sera from systemic sclerosis patients in the Stanford cohort were collected as previously published under IRB #12047(*2*). Sera from systemic sclerosis patients in the Johns Hopkins cohort were collected as previously published under NA_00039566 and IRB00226995 (*2*).

### XIST autoantigen array analysis

Fragments of XIST-associated proteins were chosen as previously described and collected patient sera were assayed by bead array as previously reported(*2*). In brief, samples were prepared by loading 25 μl of randomized patient serum or blank control onto a 96-well plate, which was subsequently shipped to SciLifeLab (Sweden) for suspension bead array assay. Non-specific reactivity was adjusted for by subtracting the MFI signal of “bare beads” from the raw MFI signals of patient samples. To compare groups with high and low reactivities, percentile ranks were used within cohorts to avoid batch effects.

### Statistical methods

All analyses were performed using R Statistical Software (v4.4.1; R Core Team 2023). Unconditional exact tests were performed using the Exact (Unconditional Exact Test) package (v3.3; (*16*)) using *exact.test* with the CSM (Convexity, Symmetry, and Minimization) method.

## Supporting information

Table S1: Number of autoantigens for autoimmune disease in AAgAtlas and female to male ratio

## List of Supplementary Materials

Table S1. Number of autoantigens for autoimmune disease in AAgAtlas and female to male ratio (related to Fig 1E).

## Acknowledgments

We thank Dr. Mark M. Davis (Stanford) and Dr. P.J. Utz (Stanford) for helpful discussion. We also thank Dr. Ji Soo Kim (Johns Hopkins) for statistical guidance.

## Funding

Howard Hughes Medical Institute (HYC)

Scleroderma Research Foundation (HYC, JL, BY)

Stanford University School of Medicine Dean’s Fellowship (BY)

Dermatology Foundation (JL)

NIH/NIAMS K24 AR080217, R01 AR073208, and P30 AR070254 (AAS)

Donald B. and Dorothy L. Stabler Foundation and the Sara and Alex Othon Research Fund (AW, GL, AAS)

Chresanthe Staurulakis Memorial Fund (Johns Hopkins Scleroderma Center Research Registry)

## Author contributions

Conceptualization: HYC

Methodology: BY, JL, YZ

Investigation: BY, JL, SS, QS, DRD, RER, SD, SN, AW, GL, YZ, MS, HRT, AAS, DFF, LSC, HYC

Visualization: BY, JL, SS

Funding acquisition: HYC

Project administration: BY, HYC

Supervision: HYC

Writing – original draft: BY, JL, HYC

Writing – review & editing: BY, JL, SS, QS, DRD, RER, YZ, AAS, LSC, HYC

## Data and materials availability

All data are available in the main text or the supplementary materials.

