## Supplementary material for "Autoantibody hotspots reveal origin and impact of immunogenic XIST ribonucleoprotein complex": Table S1: Number of autoantigens for autoimmune disease in AAgAtlas and female to male ratio

| Autoimmune diseases | Number of autoantigens in this disease | Number of autoantigens derived from XIST-asspciated proteins in this disease | p value (Hypergeometric test) | Female to male ratio | References for female to male ratio |
|---|---|---|---|---|---|
| Systemic scleroderma | 170 | 8 | 0.001480069 | 9 | Xing, et al. Journal of Investigative Dermatology (2022) (1) |
| Systemic lupus erythematosus | 367 | 12 | 0.002340529 | 9 | Xing, et al. Journal of Investigative Dermatology (2022) (1) |
| Rheumatoid arthritis | 157 | 7 | 0.004033633 | 4 | Xing, et al. Journal of Investigative Dermatology (2022) (1) |
| Mixed connective tissue disease | 527 | 13 | 0.016465656 | 5 | Ungprasert, Patompong, et al. Arthritis Care & Research (2016) (17) |
| Multiple sclerosis | 572 | 12 | 0.063259013 | 2.6 | Rommer, Paulus Stefan, et al. Therapeutic Advances in Neurological Disorders (2020) (18) |
| Demyelinating disease | 577 | 12 | 0.066770372 | 0.6 | Biswal, Nihar R., et al. Cureus (2023). (19) |
| Polymyositis | 33 | 2 | 0.068650359 | 1.4 | DeVere, R., and W. G. Bradley. Brain: a journal of neurology (1975) (20) |
| Juvenile rheumatoid arthritis | 34 | 2 | 0.072343934 | 2 | Van Kerckhove, Catherine, et al. Human immunology (1988) (21) |
| Sjogren's syndrome | 84 | 3 | 0.096518498 | 9 | Xing, et al. Journal of Investigative Dermatology (2022) (1) |
| Type 1 diabetes mellitus | 244 | 6 | 0.097471178 | 1 | Xing, et al. Journal of Investigative Dermatology (2022) (1) |
| Vasculitis | 70 | 2 | 0.232974468 | 0.66667 | Watts, Richard A., et al. Nature reviews rheumatology (2022) (22) |
| Inflammatory bowel disease | 336 | 6 | 0.274807736 | 0.5 | Lungaro, Lisa, et al. Journal of Personalized Medicine (2023) (23) |
| chronic rejection of renal transplant | 2558 | 33 | 0.606038117 | 1.5 | Chen, Po-Da, et al. Journal of the Formosan Medical Association (2013) (24) |
| Crohn's disease | 297 | 3 | 0.756947294 | 0.8 | Xing, et al. Journal of Investigative Dermatology (2022) (1) |
| Glomerulonephritis | 860 | 9 | 0.820422332 | 0.5 | Kazi, Ahmad M., and Muhammad F. Hashmi. "Glomerulonephritis." (2020) (25) |
